# Responses of Muscle Volume, Strength, and Motor Unit Properties to Bed Rest and Retraining in Young Males and Females

**DOI:** 10.64898/2026.11.04.764387

**Authors:** Fabio Sarto, Miloš Kalc, Ornella Caputo, Katarina Puš, Riccardo Magris, Giovanni Martino, Clarissa M. Brusco, Martino V. Franchi, Giuseppe De Vito, Diana A. Madrid Fuentes, Francesca Ferranti, Boštjan Simunič, Rado Pišot, Marco V. Narici

## Abstract

Sex differences in motor unit (MU) properties to simulated microgravity are unknown. Young males and females completed a 21-day head-down bed rest followed by a 21-day retraining. Bed rest reduced knee extensor strength and volume without sex differences. MU firing rate decreased in both groups, more in females, alongside increased serum C-terminal agrin fragment. After retraining, these measures returned to baseline, whereas strength only partially recovered, remaining below baseline.

## Main Text

Microgravity represents a unique environmental stressor that induces profound physiological adaptations and poses major challenges to astronaut health and performance during spaceflight^1^. The neuromuscular system is particularly affected by microgravity, as astronauts rapidly develop muscle atrophy and weakness, accompanied by motor alterations^2,3^. In this context, studying the properties of human motor units (MUs), the smallest functional units of the neuromuscular system, is fundamental for understanding the neurophysiological mechanisms of these functional alterations.

Ground-based analogues of microgravity, including unilateral lower limb suspension and bed rest, provide valuable experimental models that reproduce the mechanical unloading associated with spaceflight, enabling the investigation of the mechanisms underlying physiological deconditioning on Earth^4^. Previous studies have employed advanced electromyographic techniques to investigate alterations in MU behaviour in the context of simulated microgravity^5^.

Short-term (10–14 days) unloading studies conducted in males individuals reported rapid alterations in motoneuron discharge properties across different muscle groups^6,7^, accompanied by changes in MU potential (MUP) size^8^, complexity^9^, and MU conduction velocity^10^. While the results from these short-term studies seem to point towards a specific impairment of lower-threshold MUs^7^, medium-term (21-day) unloading resulted in altered electrophysiological properties of higher-threshold MUs as well, accompanied by neuromuscular junction (NMJ) transmission instability^11^.

However, noteworthy gaps in the literature remain. First, all previous investigations on changes in MU properties with unloading were exclusively conducted in males. Considering the well-established sex differences in neuromuscular features^12^ and the increasing exposure of female astronauts to spaceflight, establishing sex-specific changes in MU properties during unloading is essential to better understand the neuromuscular adaptations to spaceflight in both male and female astronauts. Second, while countermeasures during spaceflight and simulated microgravity have been extensively investigated^2,13^, the strategies for the recovery of neuromuscular function following unloading remain less explored^9,14^.

In the present study, we employed the gold-standard simulated microgravity paradigm of head-down bed rest (HDBR) model to investigate sex differences in human MU properties in response to simulated microgravity and subsequent retraining.

This study was part of the Italian Space Agency’s “Science for Bed Rest 2025” program, conducted at the Izola General Hospital (Izola, Slovenia). Eleven young healthy males (18–35 years) and nine young healthy females (18–31 years) underwent 21 days of −6° HDBR, followed by 21 days of high-intensity interval training (HIIT) performed on a cycle ergometer. Before bed rest (BDC-1), after bed rest (second day of recovery; R+2), and after the retraining (R+21), knee extensor maximal voluntary contraction (MVC) was assessed using isometric dynamometry, while vastus lateralis muscle volume was assessed by magnetic resonance imaging. Intramuscular electromyography (iEMG) recordings were obtained from the vastus lateralis to evaluate MU properties and NMJ transmission during voluntary isometric contractions at 25% MVC^9,15^. Blood samples were also collected at each time point and serum C-terminal agrin fragment (CAF) concentration was assessed by enzyme-linked immunosorbent assay (ELISA). Further information on the inclusion criteria, study design, techniques employed, and data and statistical analyses is provided in the Methods section. Study design is shown in Fig. 1A.

**Figure 1:**
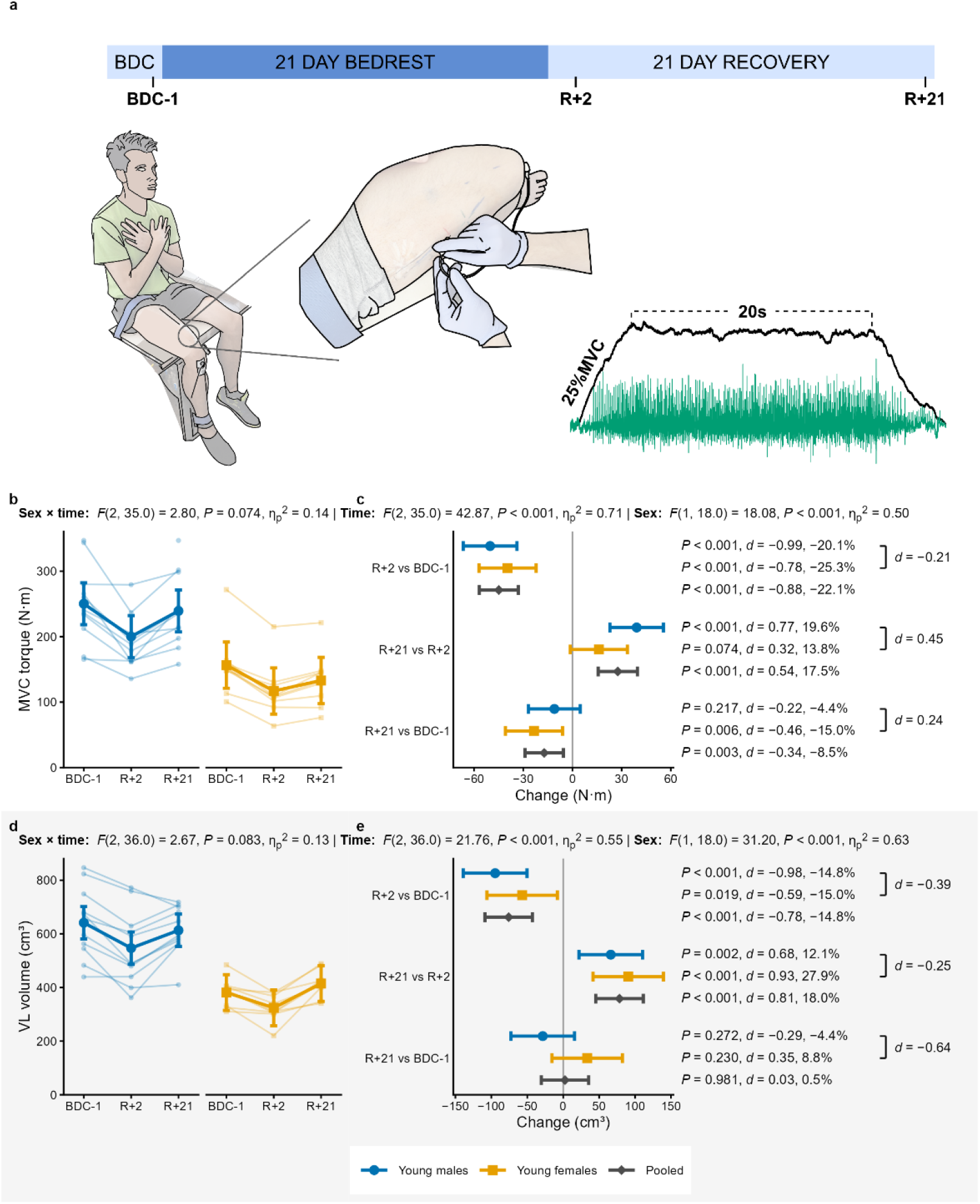
Study design and changes in knee extensor strength and vastus lateralis (VL) volume following 21 days of head-down bed rest (HDBR) and 21 days of retraining. Statistical analysis was performed using linear mixed models with sex, time and their interaction as fixed effects and participant as random intercept. Individual participant trajectories (thin lines) and estimated marginal means with 95% confidence intervals (thick lines) are shown for males and females. In the right panels, model-estimated changes between timepoints are shown within each sex and averaged over sex (Pooled), with 95% confidence intervals, exact P values adjusted for multiple comparisons, Cohen’s d and percentage change; brackets report d for the sex difference in each change (males minus females). Study timeline and representative setup for intramuscular electromyography (iEMG) recording during a trapezoidal contraction at 25% of maximal voluntary contraction (MVC), with the force trace (black) and the iEMG signal (green) (a); MVC torque (b) and its changes (c); VL volume (d) and its changes (e). BDC-1: last baseline day; R+2: second day of recovery; R+21: end of recovery.

Knee extensor MVC declined similarly in males and females following HDBR (−22.1% averaged over sex, P < 0.001; Fig. 1B-C). Although MVC increased in both groups at R+21 compared with R+2, it remained 8.5% below baseline (P = 0.003). Vastus lateralis volume was similarly reduced in males and females at R+2 (−14.8% averaged over sex, P < 0.001; Fig. 1D-E) and fully recovered in both groups. The declines in muscle strength and volume with HDBR are consistent with estimates from meta-analyses^16^ and confirm the effectiveness of the unloading model employed. Our findings of comparable relative muscle atrophy and weakness in males and females add to the limited and inconsistent literature on sex differences in response to simulated microgravity^17–20^. In addition, to our knowledge, this is the first study to compare males and females with a HIIT-based recovery following simulated microgravity. Together, these results suggest that this intervention was accompanied by full recovery of muscle volume but only partial recovery of muscle strength. We have previously reported complete recovery of muscle strength and size with resistance training-based recovery in young healthy males following unilateral lower limb suspension^21,22^, while incomplete strength recovery was observed with endurance-based retraining following horizontal bed rest^14^. This may suggest that retraining modality plays a role in the kinetics of muscle strength recovery following unloading.

To identify potential neurophysiological determinants of these functional changes, we assessed MU properties by iEMG. The number of MUs analysed is reported in Table 1.

**Table 1:**
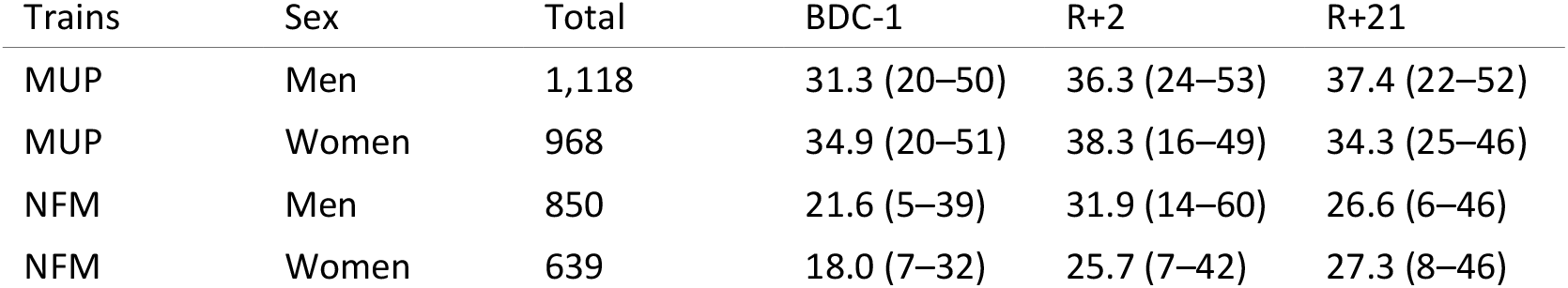
Number of motor units analysed. MUP, motor unit potential trains (MUP area and turns); NFM, near-fibre MUP trains (NFM jiggle). Total number of trains over the three timepoints; BDC-1, R+2 and R+21, mean (range) number of trains per participant.

| Trains | Sex | Total | BDC-1 | R+2 | R+21 |
|---|---|---|---|---|---|
| MUP | Men | 1,118 | 31.3 (20–50) | 36.3 (24–53) | 37.4 (22–52) |
| MUP | Women | 968 | 34.9 (20–51) | 38.3 (16–49) | 34.3 (25–46) |
| NFM | Men | 850 | 21.6 (5–39) | 31.9 (14–60) | 26.6 (6–46) |
| NFM | Women | 639 | 18.0 (7–32) | 25.7 (7–42) | 27.3 (8–46) |

Motoneuron firing rate decreased by 8.7% after bed rest (P = 0.002; Fig. 2A, B), increased by 13.3% after retraining (P < 0.001), and did not differ from baseline at R+21. The changes across bed rest and retraining were numerically larger in females than in males: firing rate decreased by 10.7% in females (d = −0.50) and by 6.6% in males (d = −0.29) after bed rest, and increased by 18.3% (d = 0.75) and 8.3% (d = 0.34), respectively, after retraining. In females, firing rate at R+21 was slightly above baseline (+5.6%, P = 0.209), whereas in males it returned to baseline (+1.2%, P = 0.918). While our results confirmed previous observations of declining firing rates with unloading and subsequent increases with retraining^7,9,14^, female motor units may be more responsive to both unloading and retraining. However, as the sex × time interaction was not significant (P = 0.148), this hypothesis warrants cautious interpretation and further investigation in larger samples. Sex differences in neuromodulation, including persistent inward currents^23^, which we previously showed to be a key determinant of firing rate in the context of unloading and recovery^14,24^, may contribute to this behaviour.

**Figure 2:**
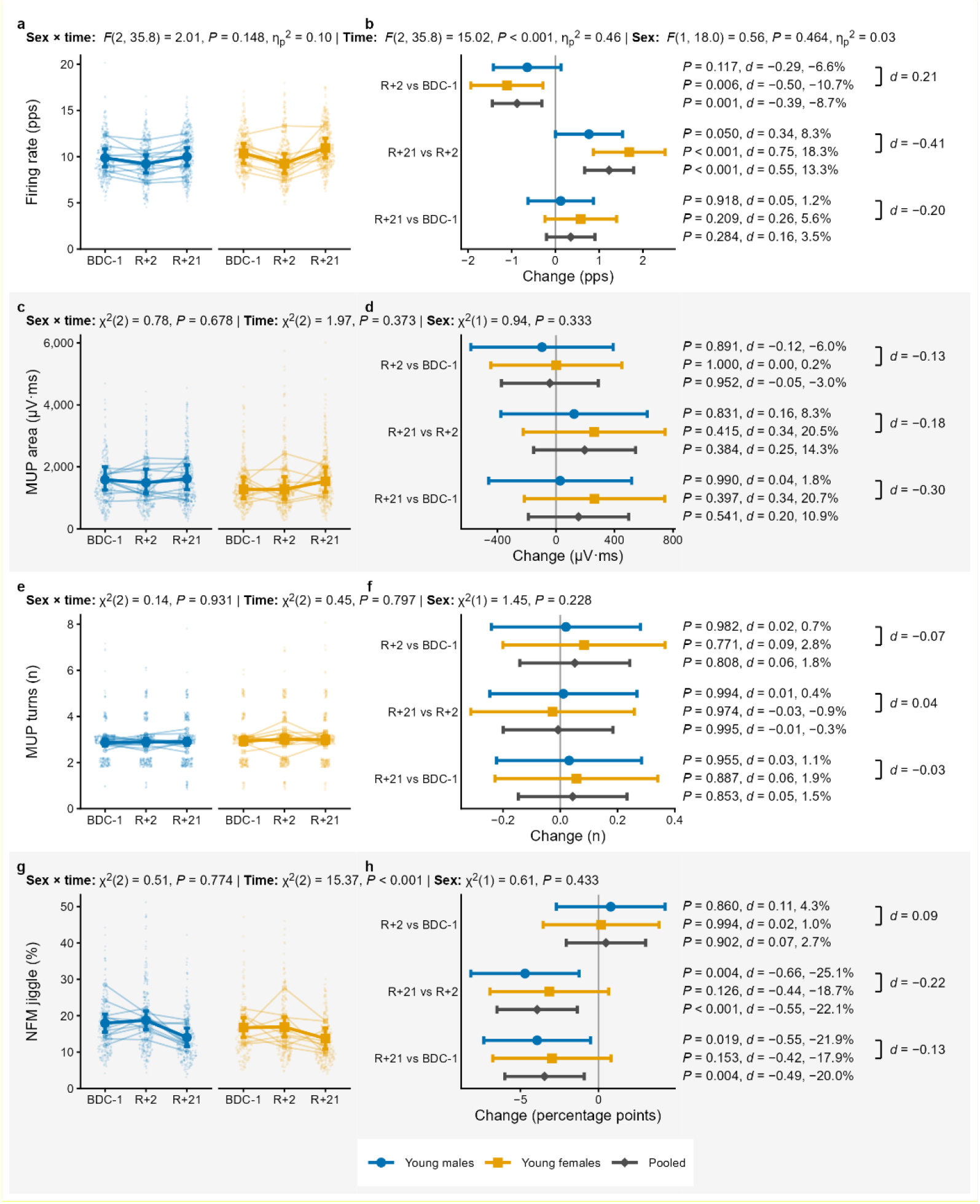
Changes in motor unit (MU) firing rate, near-fibre MU potential (NF-MUP) jiggle, MUP area and MUP turns, obtained with iEMG following 21 days of HDBR and 21 days of retraining. Statistical analysis was performed using linear mixed models (firing rate) and generalised linear mixed models (NF-MUP jiggle, MUP area, MUP turns) with sex, time and their interaction as fixed effects, and random intercepts for participant and recording session; individual MUs were the unit of observation. Small dots represent individual MUs, thin lines participant means and thick lines estimated marginal means with 95% confidence intervals, for males and females. In the right panels, model-estimated changes between timepoints are shown within each sex and averaged over sex (Pooled), with 95% confidence intervals, exact P values adjusted for multiple comparisons, Cohen’s d and percentage change; brackets report d for the sex difference in each change (males minus females). Firing rate (a) and its changes (b); NF-MUP jiggle, a proxy of neuromuscular junction transmission stability (c) and its changes (d); MUP area (e) and its changes (f); MUP turns (g) and their changes (h). pps: pulses per second. BDC-1: last baseline day; R+2: second day of recovery; R+21: end of recovery.

In contrast to firing rate, MUP area (Fig. 2C-D), reflecting MU size, and complexity (number of turns; Fig. 2E-F), associated with myofibre denervation and muscle ion channel dysfunction, were unaffected by HDBR and retraining, with no differences between sexes. These data contrast with previous findings in young males, which reported increases in MUP complexity^9,11^ and decreases in MUP area^11^, but are in line with another recent study reporting no changes in either parameter^8^ in response to unloading. Discrepancies among studies may reflect high inter-individual variability combined with the relatively small samples typical of unloading research, which is costly and logistically complex.

NMJ transmission stability, evaluated by NF-MUP jiggle, showed a significant time effect (P < 0.001; Fig. 2G, H). Jiggle did not change after HDBR but decreased by 22.1% after retraining (P = 0.001), ending 20.0% below baseline (P = 0.004). The response was similar in both sexes (sex × time, P = 0.774), with comparable reductions in males (−21.9%) and females (−17.9%) relative to baseline. Improvements in NF-MUP jiggle relative to baseline values were previously observed with resistance training-based recovery following unloading^9^, suggesting that exercise-based recovery can improve NMJ transmission following simulated microgravity.

To assess whether these electrophysiological changes were accompanied by NMJ remodelling, we measured serum CAF, a circulating biomarker of NMJ degradation (Fig. 3A, B). CAF concentration changed over time (P < 0.001), increasing by 16.2% after bed rest (P < 0.001) and returning to baseline after retraining (−12.8% vs REC2, P < 0.001; +1.4% vs BDC-1, P = 0.928). The response was comparable in males and females (sex × time, P = 0.482), with increases of 12.3% and 20.4% after bed rest, respectively. Simulated microgravity therefore appears to induce a transient NMJ stress response that resolves during retraining. This agrees with previous reports of increased CAF with unloading and, despite some inconsistencies, of generally favourable CAF responses to exercise or chronic physical activity^25^. As we previously observed^9^, CAF and jiggle changed at different times. This temporal mismatch may suggest that NMJ remodelling/instability precedes functional alterations in NMJ transmission.

**Figure 3:**
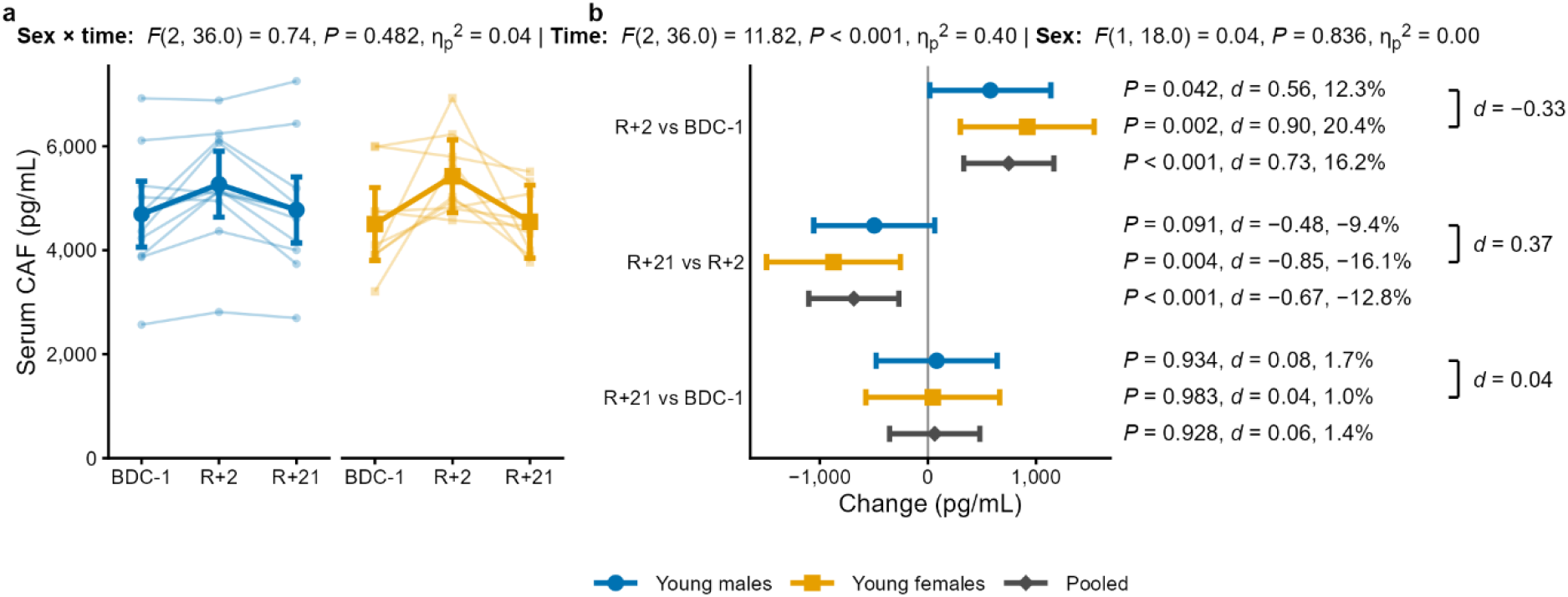
Changes in serum C-terminal agrin fragment (CAF) following 21 days of HDBR and 21 days of retraining. Statistical analysis was performed using linear mixed models with sex, time and their interaction as fixed effects and participant as random intercept. Individual participant trajectories (thin lines) and estimated marginal means with 95% confidence intervals (thick lines) are shown for males and females. In the right panel, model-estimated changes between timepoints are shown within each sex and averaged over sex (Pooled), with 95% confidence intervals, exact P values adjusted for multiple comparisons, Cohen’s d and percentage change; brackets report d for the sex difference in each change (males minus females). Serum CAF concentration (a) and its changes (b). BDC-1: last baseline day; R+2: second day of recovery; R+21: end of recovery.

In conclusion, 21 days of simulated microgravity caused comparable losses of knee extensor strength and volume in young males and females, together with a transient NMJ stress response and a reduction in motoneuron firing rate that appeared greater in females. HIIT-based retraining restored muscle volume, NMJ integrity and motoneuron firing rate, and improved NMJ transmission beyond baseline, whereas strength recovery remained incomplete, pointing to determinants beyond motoneuron discharge and NMJ transmission (e.g., muscle and tendon stiffness). These findings are relevant for understanding human neuromuscular adaptations during spaceflight and recovery and may inform the development of sex-specific countermeasures.

## Methods

### Study design

The bed rest campaign was conducted between August and October 2025 at the General Hospital Izola (Izola, Slovenia), coordinated by the Science and Research Centre Koper in collaboration with Italian research partners and under the supervision of the Italian Space Agency (ASI). It was approved by the National Medical Ethics Committee of the Republic of Slovenia (no. 0120-161/2025-2711-7) and performed in accordance with the Declaration of Helsinki. All participants were informed about the experimental procedures and provided written informed consent.

The protocol comprised 2 days of baseline data collection (BDC), 21 days bed rest (BR) and 21 days of recovery (R). Participants stayed in hospital for 26 consecutive days (BDC-2 – BDC-1, BR and R+1–R+3) and spent the remaining 18 days of recovery at home. Two participants started the protocol each day. A final one-day assessment took place in the hospital at the end of recovery scheduled between R+0 and R+24, here marked as R+21 for convenience. The measurements reported in this study were collected on the last baseline day (BDC-1), on the second day of recovery (R+2) and at the final assessment (R+21).

Of more than 300 applicants, 24 were selected and 20 young healthy adults (11 males and 9 females) were included. Participants were aged 18–35 years, had a body mass index of 22–32 kg m^−2^, passed medical and kinesiological examinations, were non-smokers and had no severe acute or chronic disease, metallic implants or participation in another biomedical study in the previous 3 months. Females participants entered the protocol immediately after the last day of menstruation. All included participants completed all experimental procedures, except for one male participant who was not able to perform the MVC test at R+2 because of knee pain, thus his MVC and iEMG data from this timepoint were excluded from the analysis.

### Head-down bed rest

Participants were housed in air-conditioned hospital rooms (7–9 per room, separated by sex) under 24-h medical care surveillance. During BR, participants were in the 6° head-down position; a pillow was allowed for head support, and one shoulder was always kept in contact with the mattress. Daily living activities, including eating, personal hygiene, urination and leisure activities, were performed in the tilted position; defecation and showering were performed in the horizontal position in the bathroom. No exercise or countermeasures were allowed, except passive stretching of the lower back. Sleep was scheduled from 22:00 to 06:00.

Throughout the hospital stay, participants received standardized, non-individualised hospital meals four times a day (breakfast, lunch, snack and dinner) with a macronutrient composition of 60% carbohydrate, 25% fat and 15% protein. Energy requirement was set at the group level as resting energy expenditure multiplied by 1.2 during BR and by 1.4 during ambulatory periods.

### Retraining

During recovery, participants completed a personalised high-intensity interval training (HIIT) on a power-controlled stationary cycle ergometer, three sessions per week for 3 weeks. The first session took place on the last day of the hospital stay (R+3), and the remaining eight sessions at a local fitness facility under the supervision of a certified trainer. Bout workloads were set between the cycling power at the gas exchange threshold and the peak aerobic cycling power measured after BR, and the load progressed from five to seven 4-min bouts per session (39–51 min per session).

### Muscle strength and intramuscular electromyography recording

Quadriceps muscle strength was assessed using an isometric dynamometer (RS 206-0290; Tedea Huntleigh, Selb, Germany) with the knee positioned at 90° of flexion. Three trials, with 1min rest, were performed and the highest force value was retained as the maximum voluntary contraction (MVC).

Motor unit potential (MUP) trains were extracted from the intramuscular electromyography (iEMG) signal recorded from the vastus lateralis (VL) muscle during submaximal isometric contractions. The VL motor point was first identified in the distal portion of the muscle (approximately 30% of femur length) using low-intensity percutaneous electrical stimulation (8– 16 mA; Digitimer Ltd., Welwyn Garden City, Hertfordshire, UK). The motor point was defined as the region eliciting the largest muscle twitch. Ultrasound imaging was then used to measure subcutaneous adipose tissue thickness and muscle width to guide the selection of the appropriate concentric needle electrode length (25 or 37 mm) for insertion at the motor point (S53155, Teca Elite, Natus Medical Inc., Middleton, WI, USA). Following needle insertion, participants performed six 30-s trapezoidal isometric contractions at 25% of MVC, while receiving real-time visual feedback of the force signal and target trapezoidal template. Each contraction consisted of a linear increase in force, a plateau at the target force, and a subsequent linear decrease at the same rate as the increase. For each contraction, the needle was rotated 180° within the same insertion site and slightly withdrawn before the subsequent pair of contractions. Trials were separated by 1 min of rest.

The iEMG signal was sampled at 40 kHz using the PowerLab acquisition toolbox and the LabChart software (v.8.13; ADInstruments, Sydney, Australia). MUP trains were extracted from the steady phase of the contraction and analysed using DQEMG software by an experienced operator (F.S.). MUP onset, end, positive peak, and negative peak markers were manually adjusted when necessary. The following MUP characteristics were assessed^9,15^: MUP area, MUP complexity via the number of turns (changes in MUP waveform of at least 20 μV) and mean firing rate. Near-fibre MUP (NF-MUP) trains were obtained by applying a second-order low-pass differentiator filter to the MUP signals and were visually inspected to exclude trains containing contaminating activity from other MUs or electric noise. We derived the NF-MUP segment jiggle, which quantifies the spatial variability between consecutive NF-MUPs and is a proxy of NMJ transmission stability.

### Magnetic Resonance Imaging

Both thighs were imaged with a 3T magnetic resonance imaging (MRI) scanner (MAGNETOM Vida Fit; Siemens Healthineers, Erlangen, Germany) using a dedicated phased array-coil (18-channel body coil and a 32-channel spine coil). Participants were scanned in a supine position. Scans were acquired after at least 8 h of head-down tilt bed rest, once body fluid shifts had stabilised; this condition was not applied at R21. The protocol included a T1 weighted (T1w) turbo spin echo sequence for high resolution anatomical reference imaging (sequence parameters: (sequence parameters: image resolution 1.1 × 1.1 × 4.0 mm3; slices: 40; no spacing between slices; matrix size: 400 × 225; repetition time (TR): 550 ms; echo time (TE): 9.80 ms; Bandwidth: 500 Hz/Px; acquisition time: 2:05 min). The protocol included an additional gradient echo Volumetric Interpolated Breath hold Examination 6pt Dixon sequence (sequence parameters: image resolution: 1.3 × 1.3 × 3.0 mm3; slices: 60, 20% spacing between slices; matrix size: 352 × 352; TR: 14.00 ms; Tes: 1.90, 3.73, 5.56, 7.39, 9.22 and 11.05 ms; Bandwidth: 710 Hz/Px; acquisition time: 1:49 min). Minimum possible TE values were selected at the given image resolution. Additional image contrasts (in-phase contrast, opposed phase contrast, water contrast and fat contrast images) were obtained from the Dixon sequence.

Thigh muscles were segmented using on MuscleMap^26^ using the water-phase images.images, and the segmentations were corrected manually in 3D Slicer (version 5.12.1). Vastus lateralis (VL) volume (cm^3^) was calculated for the whole muscle as the sum of the voxel volumes within the VL segmentation.

### Blood sampling and ELISA

Blood samples were collected at 7:00 a.m. from the medial cubital vein, allowed to clot for 30 min at room temperature, and subsequently centrifuged at 2,500 x g for 10 min at 4 °C to derive serum. Following centrifugation, serum was aliquoted and immediately stored at −80 °C until analysis. Serum samples were used for assessing the concentration of C-terminal agrin fragment (CAF), a molecular hallmark of NMJ degradation^25^, using commercially available enzyme-linked immunosorbent assay (ELISA) kits (Abcam, ab216945). Inter-plate variability was evaluated prior to determining sample concentrations by calculating the coefficient of variation (CoV) across plates. When the CoV exceeded 20%, the data were appropriately normalized using internal calibrators. For all the assays, sample concentrations were determined by interpolating absorbance values from a standard calibration curve generated using known concentrations provided with the kit. A separate standard curve was generated for each plate.

### Statistical analysis

Analyses were performed in R (version 4.5.0) with the packages lme4, lmerTest, glmmTMB, car, emmeans, effectsize and performance. Mixed models were fitted with lme410 by restricted maximum likelihood, with timepoint (BDC-1, R+2, R+21), sex and their interaction as fixed effects. For MVC and serum CAF, a random intercept for participant was included. For the iEMG parameters, individual MUs were the unit of observation. Random intercepts were included for participant and for each recording session (participant × timepoint). Model assumptions (model fit, linearity, homogeneity of variance, influential observations, collinearity and normality of residuals) were checked visually with the performance package. Normality of residuals was not met for MUP area and NFM jiggle, which were therefore analysed with generalised linear mixed models with an inverse Gaussian distribution and a log link (MUP area) and a Gamma distribution and an identity link (NFM jiggle). The number of MUP turns was underdispersed relative to a Poisson distribution and was modelled with a Conway–Maxwell–Poisson generalised linear mixed model (log link) in glmmTMB. Fixed effects were tested with type III F-tests with Kenward– Roger (MVC, CAF) or Satterthwaite (FR) degrees of freedom in the linear mixed models and with type III Wald χ2 tests in the generalised linear mixed models (MUP area, MUP turns, NFM jiggle). Pairwise comparisons between timepoints were performed with emmeans within each sex and averaged over sex, with multivariate t adjustment for multiple comparisons11. These comparisons were made for all outcomes, whether or not the sex × time interaction was significant; they describe the change within each sex, whereas a difference between sexes was inferred only from the interaction. Effect sizes are reported as Cohen’s d with 95% confidence intervals. For the linear mixed models, d was calculated as the model-estimated contrast divided by the total standard deviation of the model (participant, session and residual variability). For the generalised linear mixed models, whose variance components are not on the scale of the estimated contrasts, d was calculated as the model-estimated change on the original scale divided by the standard deviation of the observations around the fixed-effect predictions. Cohen’s d and the percentage change, calculated from the estimated marginal means, are reported for every pair of timepoints within each sex and averaged over sex; d is also reported for the sex difference in each change (change in men minus change in women). Statistical significance was set at P < 0.05.

## Data availability

The data that support the findings of this study will be made available from the corresponding author upon reasonable request.

## Acknowledgments

We thank all the participants who volunteered for this bed rest investigation. We also thank the nurses and medical team at the Izola General Hospital (Izola, Slovenia), as well as the staff of the Koper Science and Research Center (Institute for Kinesiology Research) for essential support in organizing and assisting the participants throughout the study. The study project was carried out within the framework of the collaboration agreement n. 2025-4-HB.0 between the Italian Space Agency (ASI) and the University of Padova, Department of Biomedical Sciences, for the “Neuromyo” project.

## Author contributions

F.S., B.S., R.P., M.V.F., G.D.V., and M.V.N. conceptualized and designed the study; M.V.N. obtained the funding; B.S., R.P., F.F., and M.V.N. coordinated the bed rest study; F.S., M.K, O.C., K.P., R.M., G.M., and C.M.B. performed data collection; F.S. performed iEMG analysis; O.C. performed ELISA to assess CAF; K.P. and D.A.M.F. analysed MRI data; M.K. performed statistical analysis and prepared the figures; F.S., M.K., and O.C. drafted the manuscript and all the authors revised it.

